# Activation of the Cpx-envelope stress response system promotes tolerance to antibacterials delivered by arginine-rich peptides and aminoglycosides in *Escherichia coli*

**DOI:** 10.1101/2020.08.31.274910

**Authors:** Jakob Frimodt-Møller, Andreas Koulouktsis, Godefroid Charbon, Marit Otterlei, Peter E. Nielsen, Anders Løbner-Olesen

**Author notes:** Corresponding authors: Jakob Frimodt-Møller and Anders Løbner-Olesen.

## Abstract

Cell penetrating peptides (CPP) are increasingly used for cellular drug delivery in both pro- and eukaryotic cells, and oligoarginines have attracted special attention. However; their mechanism of action, particularly for prokaryotes is still unknown. Arginine-rich CPPs (R-CPP) efficiently delivers the antimicrobial peptide nucleic acid (PNA) into bacteria. Here, we show that resistance to an R-CPP PNA conjugate in *Escherichia coli* requires multiple genetic modifications and is specific to R-CPP and not to the PNA-part. An integral part of the resistance was the constitutively activated Cpx-envelope stress response system (*cpx*\*), which decreased the cytoplasmic membrane potential and thereby indicates an indirectly energy dependent uptake mechanism. Interestingly, *cpx*\* mutants also showed increased tolerance to aminoglycosides and R-CPP conjugated to a peptide targeting the DNA sliding clamp; i.e., similar uptake in *E. coli* for these antimicrobial compounds. We speculate that the *cpx*\* phenotype could create an evolutionary opportunity to adapt and evolve in the presence of either compounds.

**Author summary:** The emergence of multidrug resistant bacteria is raising the need for new classes of antibiotics. Peptide nucleic acids (PNAs) may fill this requirement by their ability to block translation of essential mRNAs and hence inhibit growth. PNA needs conjugation to a delivery peptide (cell penetrating peptide; CPP) to enter the bacteria. Arginine-rich CPPs (CPP_R_) are receiving a lot of attention for use as delivery vessels. Here, we show, for the first time, CPP_R_-PNA resistance in *Escherichia coli* directed towards the delivery peptide. Consequently, resistance also applies to other antimicrobial compounds delivered by the same carrier. An integral part of CPP_R_ resistance is due to a constitutive active Cpx-response system, which leads to a decreased electric potential (Δ*Ψ*) across the inner membrane. The decreased Δ*Ψ* is a result of down-regulation of two aerobic respiratory operons, namely NADH:ubiquinone oxidoreductase complex I and cytochrome bo_3_ ubiquinol oxidase. The decreased Δ*Ψ* also led to increased tolerance to aminoglycosides. This shows that a (large) negative Δ*Ψ* is important for providing sufficient free energy for membrane translocation of both CPP_R_ and that the inner membrane is the main barrier for entry of both arginine-rich delivery peptides and aminoglycosides.

## Introduction

Antimicrobial resistance is one of the major challenges of the 21st century. Thus, efforts to bring novel antimicrobial compounds, including antimicrobial peptides (AMP), into clinical use are accelerating. Antisense technology, as a gene targeted precision drug modality has recently produced several new drugs in clinical use (Inotersen, Valonesorsen and Golodirsen), and agents based on the pseudopeptide DNA mimic, peptide nucleic acid (PNA)(1) and phosphorodiamidate morpholino oligonucleotide (PMO)(2) have been proposed as a future antibiotic. Naked PNA and PMO, like many AMPs, are unable to traverse the cell envelope in *Escherichia coli*(2, 3). Hence, these requires a carrier molecule (i.e., cell-penetrating peptide (CPP)) for efficient translocation into the cytoplasm. Very little is known about how CPP conjugated to PNA/PMO/AMP enters the cell and for CPP-PNA only one paper has addressed this topic(4). Here, PNA is designed to target the translation initiation region of the mRNA of an essential gene causing steric hindrance of ribosome binding, and thereby inhibition of gene expression and cell death(1). PNA conjugated to the lysine-rich CPP *L*((KFF)_3_K) (CPP_KFF_-PNA) enters *E. coli* by the non-essential, inner membrane transporter SbmA(4); consequently SbmA deficient *E. coli* are highly resistant to this peptide-PNA. However, PNA conjugated to an arginine-rich CPP, (R-Ahx-R)_4_-Ahx-(βAla)-PNA (CPP_RXR_-PNA) enters the cell in an unknown and SbmA-independent manner(4). Cationic arginine-rich CPPs (R-CPP) are increasingly used as molecular carriers and several CPP-based drugs are currently being pursued in drug discovery in relation to for example wound healing, cancer, and Duchenne Muscular Dystrophy(5). How R-CPP enters eukaryotic and prokaryotic cells is poorly understood. In eukaryotes, evidence supports predominantly an endocytotic (energy-dependent) pathway, although direct translocation (energy-independent mechanism) may also play a role (5), while it is unclear how R-CPPs penetrate through membrane lipids in prokaryotes.

The envelope of Gram-negative bacteria consists of three essential structurally and chemically diverse layers: 1) the inner membrane (cytoplasmic membrane); 2) the periplasm containing a thin peptidoglycan layer; and 3) the outer membrane that contains phospholipids in the inner leaflet and lipopolysaccharide (LPS) in the outer leaflet. *E. coli* can actively transport compounds across the cytoplasmic membrane using either ATP directly via membrane transporter proteins or indirectly via the membrane potential (partly created by the chemical proton gradient, and also being part of theproton motive force (PMF), generated by the electron transport chain). The energetics of the cell varies considerably between growth with or without oxygen, with oxidative growth conditions providing the highest energy turnover. For instance, PMF is decreased during anoxic growth conditions. The electron transport chain will under aerobic conditions predominantly contain NADH:ubiquinone oxidoreductase complex I (organized from the *nuoABCEFGHIJKLMN*-operon; NDH-I) and cytochrome *bo*_3_ ubiquinol oxidase (organized from the *cyoABCDE* operon; cytochrome *bo*_3_ oxidase)(6). PMF generated by translocating protons from the cytoplasm into the periplasmic space(6), serves as energy storage, which is used to drive ex. ATP synthase, efflux pumps, and transport of metabolites(7). ATP is generated from ADP and inorganic phosphate by the multi-subunit protein complex F_1_F_0_-ATPase(8). F_1_F_0_-ATPase consist of two structural domains; F_1_ containing the catalytic core and F_0_ containing the membrane proton channel(8). The PMF across the cytoplasmic membrane is a combination of the electric potential (Δ*Ψ*) and the transmembrane proton gradient (Δ*pH:* internal pH – external pH). Δ*Ψ* is always negative inside growing neutrophils, such as *E. coli*, and Δ*Ψ* decreases as Δ*pH* increases(9); i.e. bacteria may regulate Δ*Ψ* and/or Δ*pH* in order to control PMF.

The Bae, Cpx, Psp, Rcs, and σ^E^ pathways constitute the *E. coli* signaling systems that detect and respond to alterations of the bacterial envelope(10). These pathways are involved in the biogenesis, maintenance, and repair of the bacterial envelope, and thus contribute to cell surface integrity(10). One of the most well studied extracytoplasmic stress response-systems is the CpxA/R two-component signal transduction system, which among other genes down-regulate respiratory operons when activated(11). Like many other histidine kinases, CpxA is localized to the cytoplasmic membrane through two transmembrane helices and contains both periplasmic and cytoplasmic domains(12). The cytoplasmic response regulator, *cpxR*, is predicted to encode an OmpR-like transcriptional activator(12). CpxR consist of an N-terminal receiver domain (phospho-acceptor domain) and C-terminal DNA-binding domain(13). Under non-stress conditions the auxiliary regulator CpxP inhibits the Cpx-response by a predicted direct interaction with the periplasmic domain of CpxA(14). Under Cpx-inducing conditions, CpxP is degraded by the periplasmic protease and chaperone DegP(15). The CpxP relieved CpxA autophosphorylates on a conserved histidine residue (His-248), using ATP as its phosphoryl donor(16). Subsequently, phosphorylated CpxA donates its phosphoryl group to the conserved aspartate residue (Asp-51) in the N-terminal receiver domain of CpxR. Phosphorylated CpxR binds to DNA and act as a transcriptional regulator(16). In addition, CpxA dephosphorylates CpxR-P, which ensures that CpxR remains inactive in the absence of an activating signal(17). The outer membrane lipoprotein NlpE that sense surface adhesion, acts as a ‘sentry protein’ against envelope stress and activates the Cpx-response system through an undefined interaction with CpxA(17).

Here we show that *E. coli* can develop tolerance to antisense PNA conjugated to R-CPPs through a constitutive Cpx-response that result in a decreased Δ*Ψ* across the cytoplasmic membrane. The decreased Δ*Ψ* also confers cross-resistance to aminoglycosides indicating a similar uptake mechanism for the two classes of compounds.

## Results

To understand bacterial uptake and resistance to PNA R-CPP conjugates, we studied the widely used CPP_RXR_(4) (Table 1). Unless stated otherwise the PNA part always targets translation of the *acpP* mRNA, encoding the acyl carrier protein, which is essential for fatty acid synthesis(4). The CPP_RXR_-PNA exhibited a minimum inhibitory concentration (MIC) of 0.5 μM towards *E. coli*, whereas the corresponding CPP_RXR_-PNA containing two mismatches in the PNA sequence (PNA_mm_) did not inhibit bacterial growth up to 16 μM (Table 1); *i.e*., the CPP_RXR_ part alone did not confer any antimicrobial effect supporting an antisense mechanism of action. As a previous study failed to identify single-gene deletion mutants of *E. coli* resistant to CPP_RXR_-PNA(4), we used adaptive laboratory evolution (ALE; see Methods) to generate CPP_RXR_-PNA resistant mutants. Wild-type cells were grown in successively increasing concentrations of CPP_RXR_-PNA, to select for mutation(s) that accumulate during long-term exposure to CPP_RXR_-PNA. The ALE experiments were done in a biological duplicate (two independent linages) and was terminated when resistant clones appeared after 20 successive passages, corresponding to a 16-fold increase in MIC (from 0.5 μM to 8 μM) for CPP_RXR_-PNA (Table 1).

**Table 1.**
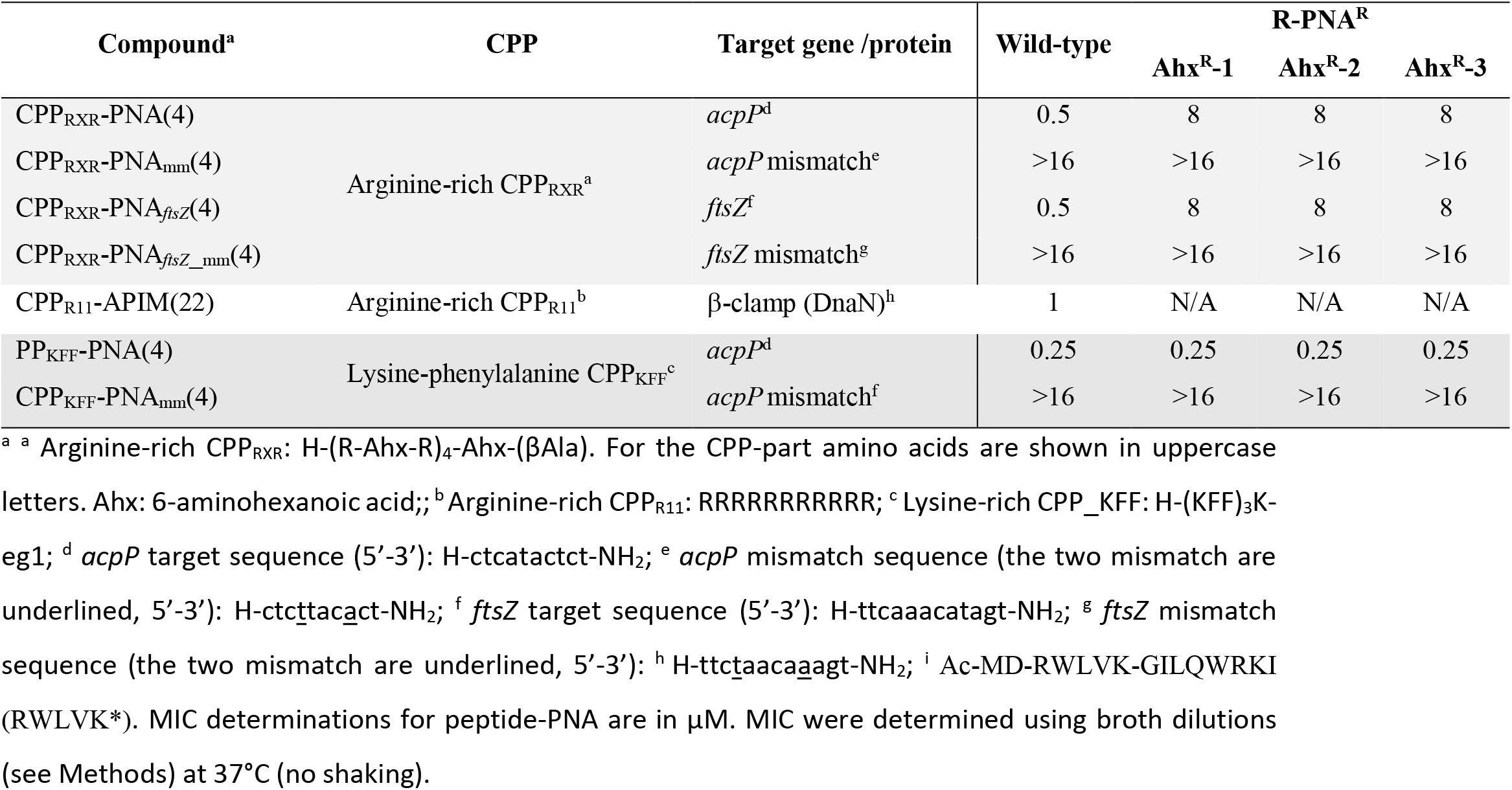
Minimum inhibitory concentration to peptide-PNA (in μM).

Five individual clones from each experiment at day 20 (ten clones in total) were isolated and sequenced (Table 2; Table S1; Table S2). From the ten CPP_RXR_-PNA resistant clones we identified three different genotypes, which were annotated as such: Ahx^R^-1 to 3. Ahx^R^-1 and 2 were isolated from lineage one and Ahx^R^-3 from lineage two. None of the mutations identified in the CPP_RXR_-PNA resistant clones resulted from adaptation to the growth medium, as none of them occurred in wild-type cells adapted to the same growth medium without CPP_RXR_-PNA (Table S3).

**Table 2.**
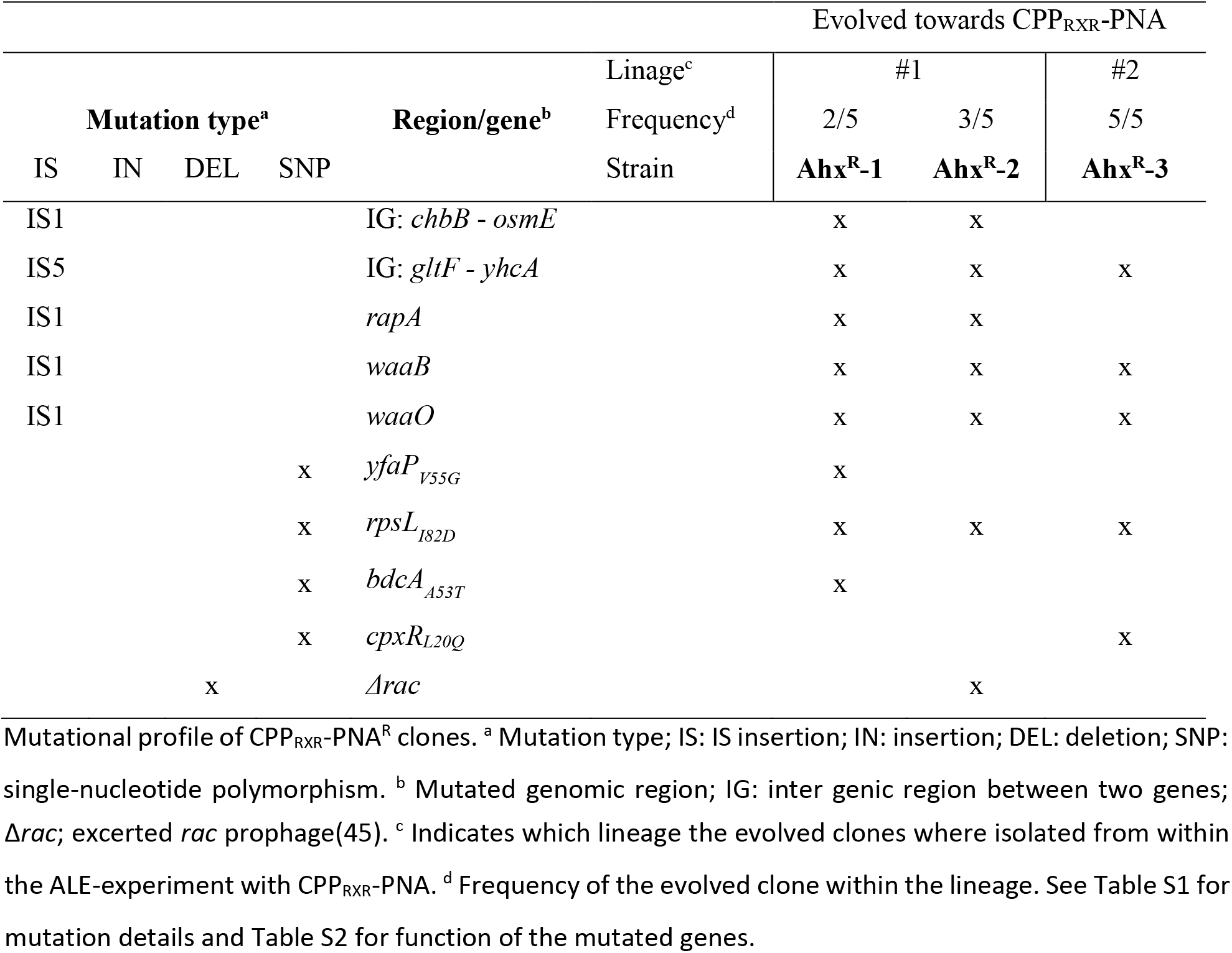
CPP_RXR_-PNA resistant genotypes.

Linage two from the ALE-experiment appeared to be clonal. This lineage contained genes/operons that were mutated in all of Ahx^R^-1 to 3; the *gltF-yhcA* intergenic region, *waaB, waaO*, and *rpsL*, as well as a mutation in *cpxR* specific to only Ahx^R^-3 (Table 2). *gltF* belongs to the *gltBDF*-operon, where *gltB* and *gltD* encode the large and small subunit of glutamate synthase(18), respectively. The function of GltF and the hypothetical protein YhcA, is unknown. The *waa*-operon contains the major core-oligosaccharide assembly operons in *E. coli(19*). RpsL is the only mutated gene that is essential. The product of *rpsL* is the S12 protein; a component of the 30S subunit of the ribosome where it plays a role in translational accuracy(20). Lastly, *cpxAR* encodes the two-component regulatory system CpxA/R(21) (see above).

### Resistance to CPP_RXR_-PNA is towards the arginine-rich CPP-part and not to PNA-part

The MIC were determined for Ahx^R^-1-3 (collectively CPP_RXR_-PNA^R^) towards additional peptide-PNA conjugates (Table 1). R-PNA^R^ strains were also resistant to the analogous CPP_RXR_ targeting the essential division protein *ftsZ* instead of *acpP:* CPP_RXR_-PNA_*ftsz*_ (Table 1). On the contrary, all R-PNA^R^ strains remained sensitive to an *acpP* targeting PNA conjugated to the lysine-phenylalanine CPP, (KFF)_3_K (CPP_KFF_-PNA) (Table 1). Together this shows that resistance to CPP_RXR_-PNA is not directed towards the PNA-part but towards the (arginine-rich) delivery peptide, supporting the notion of different uptake mechanism for conjugates with these two peptides.

### An activated Cpx-response leads to R-CPP tolerance

The Cpx-system is known to be involved in extracytoplasmic stress response and was mutated in five out of ten sequenced CPP_RXR_-PNA resistance clones. Thus, we proceed to investigate the role of the Cpx-response in tolerance to R-CPP. The highest tolerance level to CPP_RXR_-PNA was 8 μM (i.e. CPP_RXR_-PNA^R^ level), which we define as CPP_RXR_-PNA resistance. Therefore, everything between wild-type CPP_RXR_-PNA sensitivity (>0.5 μM) and CPP_RXR_-PNA resistance is referred to as CPP_RXR_-PNA tolerance.

Loss of the Cpx-response, by deletion of *cpxA* or *cpxR* in otherwise wild-type cells had no effect on the tolerance to CPP_RXR_-PNA (Fig. 1A). Interestingly, increased levels of CpxR led to a small but reproducible increase in tolerance to CPP_RXR_-PNA (Fig. 1A), which could indicate that Cpx activation promotes tolerance to CPP_RXR_-PNA. We proceeded to introduce the *cpxR_L20Q_* mutation identified in the ALE experiment (Table 2) into wild-type cells and included a *cpxA_Δ16-17_* mutant into this study from our strain collection (isolated in a different project). In cpxR_L20Q_, leucine is replaced by glutamine at position 20, which is part of a highly conserved hydrophobic core in the N-terminal receiver domain shared by OmpR-family proteins(13). In CpxA_Δ16-17_ threonine at position 16 and leucine at position 17 of the first CpxA transmembrane domain are missing. Both mutations increased the MIC for CPP_RXR_-PNA from 0.5 μM to 4 μM, relative to wild-type cells (Fig. 1A). Thus, although more tolerant to CPP_RXR_-PNA than wild-type cells, the *cpxR_L20Q_* and *cpxA_Δ16-17_* mutants did not reach the CPP_RXR_-PNA resistance level of the evolved mutant Ahx^R^-3. Next, the Cpx-response was tested against an oligoarginine conjugate to a peptide targeting the DNA sliding clamp (DnaB; β-clamp) (RWLVK-GILQWRKI-RRRRRRRRRRR; CPP_R11_-APIM), which has been shown to inhibit bacterial growth at micromolar concentrations both *in vitro* and *in vivo*(22) (Table 1). Loss of the Cpx-response by deletion of *cpxA* in an otherwise wild-type cell had no effect on tolerance to CPP_R11_-APIM, however tolerance was reproductively increased from 1 μM to 2 μM in the *cpxR_L20Q_* and *cpxA_Δ16-17_* mutants, relative to wild-type cells (Fig. 1B). Thus this peptide also exhibits tolerance to the Cpx-mutants but only at a 2 fold increase compared to 8 fold for the CPP_RXR_-PNA.

**Fig. 1.**
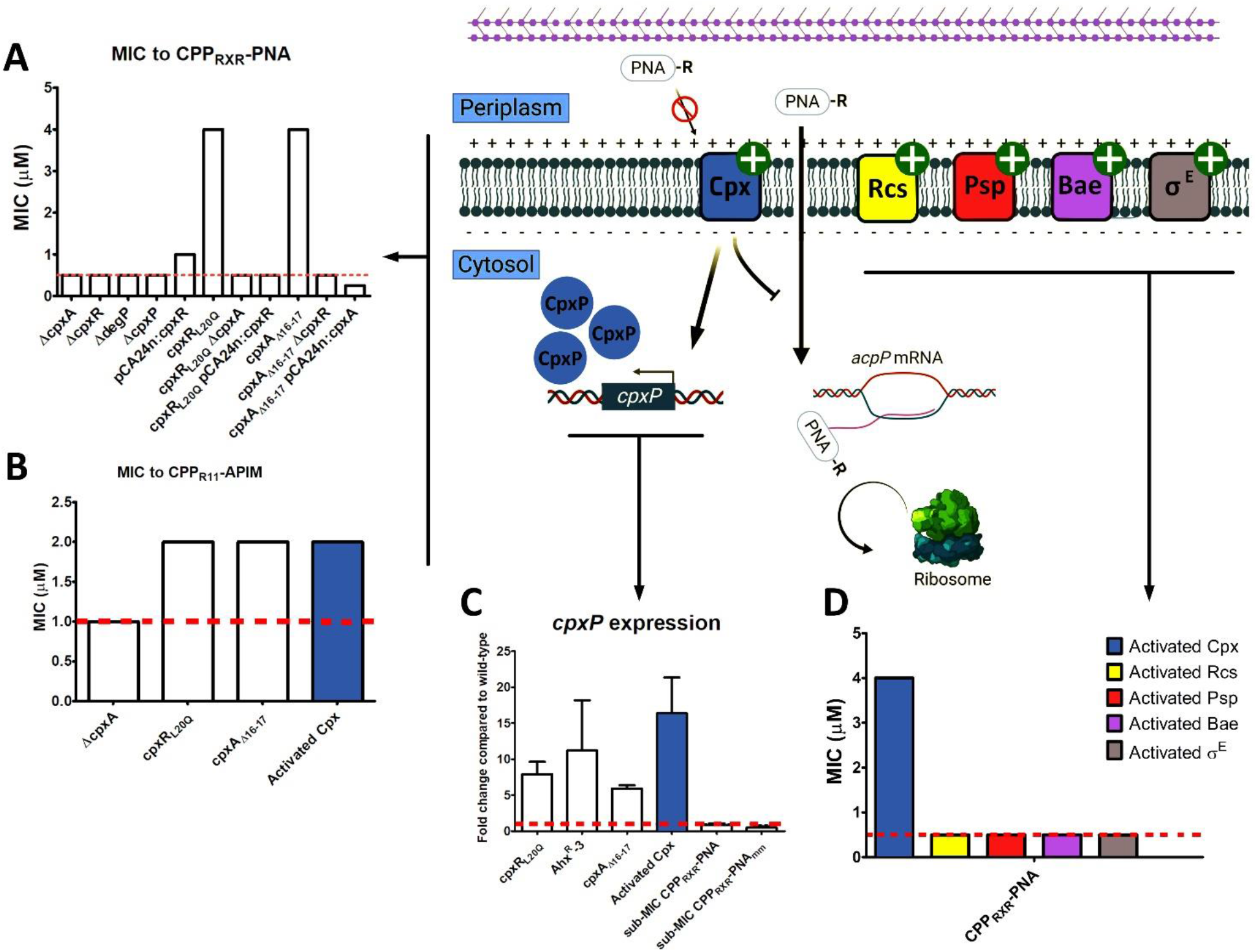
*cpxR_L20Q_* and *cpxA_Δ16-17_* activates the Cpx-response. A schematic model of the Cpx, Rcs, Psp, Bae, and σ^E^ stress response to R-PNA in the cytoplasmic membrane. Here, R-PNA does not activate the Cpx-response; however, an activated Cpx-response is the only of the five-tested extracytoplasmic stress responses that diminish R-PNA activity. The green plus sign indicates a constitutive active ESR where an activated Cpx-response confers resistance to R-PNA. A and B: MIC determinations for CPP-PNA/APIM for wild-type and mutants in μM. MIC were determined using broth dilutions (see Methods) at 37°C (no shaking). Peptide-PNA sequences are listed in Table 1. pCA24n-based plasmids were induced with 0.1 mM IPTG. C: *cpxP* expression determined by RT-qPCR (see Methods). All values are indicated as fold change relative to wild-type (dashed line). Experiments were performed in triplicate. D: MIC determinations for activated extracytoplasmic stress response-systems in μM (see text for details). The five extracytoplasmic stress responses were individually activated by overproduction of *nlpE* for the Cpx-pathway, *baeR* for the Bae-pathway, *rcsB* for the Rcs-pathway, *pspF* for the Psp-pathway, and *rpoE* for the σ^E^-pathway(10). A dashed line indicates the wild-type MIC. Peptide-PNA sequences are listed in Table 1. pCA24n-based plasmids were induced with 0.1 mM IPTG.

We used *cpxP* transcription as a readout for activation of the *cpx* response(23). *cpxR_L20Q_, cpxA_Δ16-17_*, and Ahx^R^-3 all had increased expression of *cpxP* compared to wild-type (Fig. 1C), albeit not to the level achieved by overproduction of NlpE (full activation of the Cpx-response; Fig. 1C). This confirmed that the Cpx-response was indeed activated by the *cpxR_L20Q_* and *cpxA_Δ16-17_* mutations and that these provided CPP_RXR_-PNA (and CPP_R11_-APIM tolerance) to the same level as full activation of the Cpx-response by NlpE overproduction (Fig. 1B, D). Both mutations are therefore gain-of-function mutations leading to a constitutively active Cpx-response, which previously have been denoted *cpx*\*(16, 17).

### Both *cpxR_L20Q_* and *cpxA_Δ16-17_* mutations are dependent on a functional Cpx-system for conferring CPP_RXR_-PNA tolerance

To assess whether the *cpxA_Δ16-17_* and *cpxR_L20Q_* mutations were dependent on a functional Cpx system for conferring CPP_RXR_-PNA tolerance, *cpxA* was deleted in the *cpxR_L20Q_* strain and *cpxR* was deleted in the *cpxA_Δ16-17_* strain. In either case, CPP_RXR_-PNA tolerance was reduced to wild-type level (Fig. 1A). Thus, the substitution to glutamine in position 20 in CpxR does not make it constitutively active by mimicking phosphorylation of the conserved Asp-51, and cpxR_L20Q_ still relies on a functional CpxA to be activated. Overproduction of wild-type CpxR or CpxA in the *cpxR_L20Q_* or *cpxA_Δ16-17_* mutants, respectively, also restored CPP_RXR_-PNA susceptibility to wild-type level (Fig. 1A). Together, this shows that the specific *cpxR_L20Q_* or *cpxA_Δ16-17_* mutations mediates CPP_RXR_-PNA tolerance, but do need a functional Cpx-system.

### The Cpx-response is the only extracytoplasmic stress response conferring tolerance to CPP_RXR_-PNA

The Cpx-response is only one of five extracytoplasmic stress response systems known to maintain cell envelope integrity during stress. While the Cpx-response can be activated by overproducing *nlpE*, the four other extracytoplasmic stress responses can be individually activated by overproduction of *baeR* for the Bae-pathway, *rcsB* for the Rcs-pathway, *pspF* for the Psp-pathway, and *rpoE* for the σ^E^-pathway(10). Activation of neither the Bae, Psp, Rcs, nor the σ^E^-responses led to a CPP_RXR_-PNA tolerant phenotype (Fig. 1D), showing CPP_RXR_-PNA tolerance is specific to an activated Cpx-response. However, wild-type treated with sub-MIC concentrations (0.5 X MIC) of CPP_RXR_-PNA or CPP_RXR_-PNA_mm_ did not result in an increased *cpxP* expression (Fig. 1D). Hence, although an activated Cpx-response confers CPP_RXR_-PNA tolerance, low-level CPP_RXR_-PNA treatment in itself does not trigger extracytoplasmic stress.

### Downregulation of respiratory operons leads to CPP_RXR_-PNA tolerance

Because Cpx mediates downregulation of aerobic respiratory operons(11), expression of the *nuoABCEFGHIJKLMN* and *cyoABCDE* operons were determined by RT-qPCR. As expected, transcription of *nuoA* (to evaluate expression of the *nuoABCEFGHIJKLMN* operon) and *cyoA* (to evaluate expression of the *cyoABCDE* operon) were downregulated in wild-type/pCA24n:*nlpE*, *cpxR_L20Q_, cpxA_Δ16-17_*, and Ahx^R^-3 cells (Fig. 2A). We found that such Cpx activated cells in addition to being CPP_R11_-APIM (*cpxR_L20Q_* and *cpxA_Δ16-17_;* Fig. 2B) and CPP_RXR_-PNA tolerant (Fig. 2B) were also more tolerant to the aminoglycosides gentamicin, amikacin, and kanamycin (Fig. 2C) than the wild-type. When the entire *nuoABCEFGHIJKLMN* and *cyoABCDE* operon were individually deleted from wild-type cells, the resultant Δ*nuo* and Δ*cyo* strains were more tolerant to both CPP_R11_-APIM (only tested with Δ*cyo*), CPP_RXR_-PNA and aminoglycosides relative to the wild-type (Fig. 2B,C). This confirms that CPP_R11_-APIM, CPP_RXR_-PNA, and aminoglycoside tolerance in *cpxR_L20Q_* and *cpxA_Δ16-17_* is related to aerobic oxidative phosphorylation and result from downregulation of NDH-I and cytochrome *bo*_3_ oxidase.

**Fig 2.**
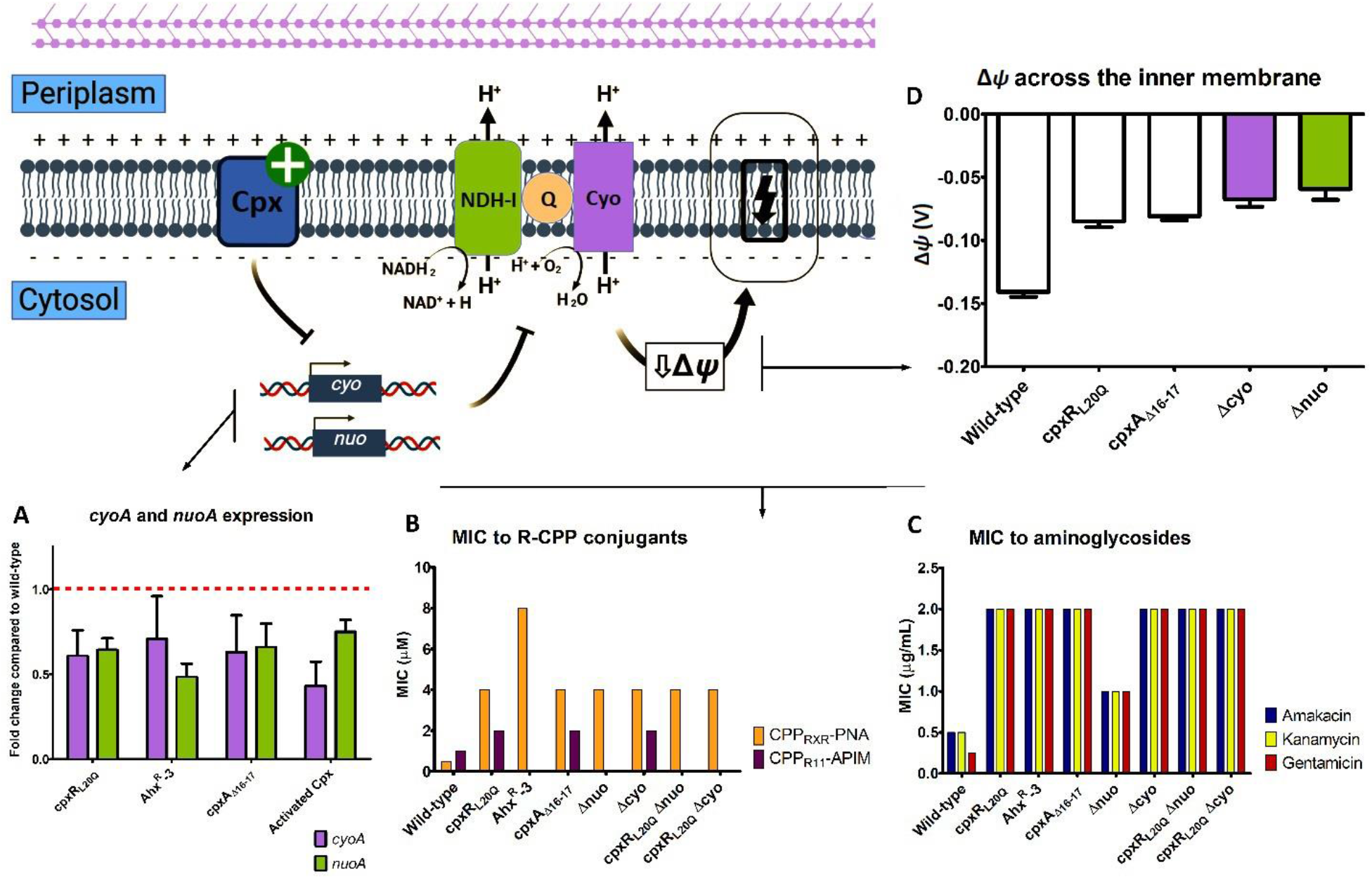
Cpx-mediated downregulation of NDH-I and cytochrome *bo*_3_ oxidase leads to a decreased Δ*Ψ* across the cytoplasmic membrane that confers resistance to CPP_R11_-APIM, CPP_RXR_-PNA, and aminoglycosides. A schematic model of an activated Cpx-response (shown here with a green plus sign) that leads to downregulation of both *nuoA* (NDH-I) and *cyoA* (cytochrome *b*o_3_ oxidase), which consequently results in a decreased Δ*Ψ* across the cytoplasmic membrane that confers cross resistance to CPP_RXR_-PNA and aminoglycosides. A: *nuoA* and *cyoA* expression determined by RT-qPCR (see Methods). All values are indicated as fold change relative to wild-type (dashed line). Experiments were performed in biological triplicates. B and C: MIC values for CPP-PNA/APIM (in μM) and aminoglycosides (in μg/mL) for wild-type and mutants. MIC were determined using broth dilutions (see Methods) at 37°C (no shaking). Peptide-PNA sequences are listed in Table 1. D: Δ*Ψ* across the cytoplasmic membrane as measured by the TPP^+^ uptake for wild-type, *cpxR_L20Q_, cpxA_Δ16-17_*, Δ*cyo*, and Δ*nuo*. Experiments were performed as a minimum in biological triplicates.

### The cytoplasmic membrane potential (Δ*Ψ*) directly affects CPP_RXR_-PNA and CPP_R11_-APIM activity

Downregulation of respiratory operons suggested that the Δ*Ψ* formed across the inner membrane plays a significant role in CPP_R11_-APIM CPP_RXR_-PNA tolerance. We determined the Δ*Ψ* at the intracellular pH 7.6 of *E. coli* based on the distribution of the lipophilic tetraphenylphosphonium ion (TPP^+^) using a TPP^+^-selective electrode(24). At this pH value the Δ*pH* is zero and the PMF is only dependent on Δ*Ψ*. Wild-type cells had a Δ*Ψ* of approx. −140 mV, in accordance with previous observations(25) (Fig. 2D). The Δ*Ψ* was reduced by the *cpxR_L20Q_* as well as the *cpxA_Δ16-17_* mutation (Fig. 2D). Δ*cyo* and Δ*nuo* cells also had a decreased Δ*Ψ* compared to wild-type; with values comparable to *cpxR_L20Q_* and *cpxA_Δ16-17_* (Fig. 2D). Thus, it is conceivable that the decreased Δ*Ψ* of *cpxR_L20Q_* and *cpxA_Δ16-17_* cells may result from downregulation of the respiratory operons.

*E. coli* maintains a stable PMF across the inner membrane, such that Δ*Ψ* decreases with an increase in Δ*pH* in an acidic medium, vice versa in an alkaline media (9) (Fig. 3). Thus, Δ*Ψ* can be modulated by growing the cells at various pH levels. In acidic media (pH 6.0) where Δ*Ψ* is decreased(9) (Fig. 3A), wild-type cells were tolerant to CPP_RXR_-PNA, CPP_R11_-APIM and gentamicin (Fig. 3A), while an increased Δ*Ψ* due to growth in alkaline media (pH 8.0)(9) resulted in sensitivity to both CPP_RXR_-PNA, CPP_R11_-APIM and gentamicin (Fig. 3B). The changes in Δ*Ψ* in the two different pH-adjusted settings had no effect on tolerance to CPP_KFF_-PNA (Fig. 3A,B). *E. coli* cells deficient of the F_1_F_0_-ATPase or supplemented with alanine have been shown to have an increase in Δ*Ψ*(26, 27) and were more sensitive to CPP_RXR_-PNA, gentamicin, and CPP_R11_-APIM (only tested with alanine), but sensitivity to CPP_KFF_-PNA was similar to wild-type (Fig. 3B). Finally, Δ*Ψ* is decreased under anaerobic growth conditions(26) explaining why wild-type under these conditions are more tolerant to CPP_RXR_-PNA and gentamicin (Fig. 3A). Most likely the effect of Δ*Ψ* on antimicrobial activity is explained by increased inner membrane passage of the polycationic molecules energetically driven by the negative membrane potential.

**Fig. 3.**
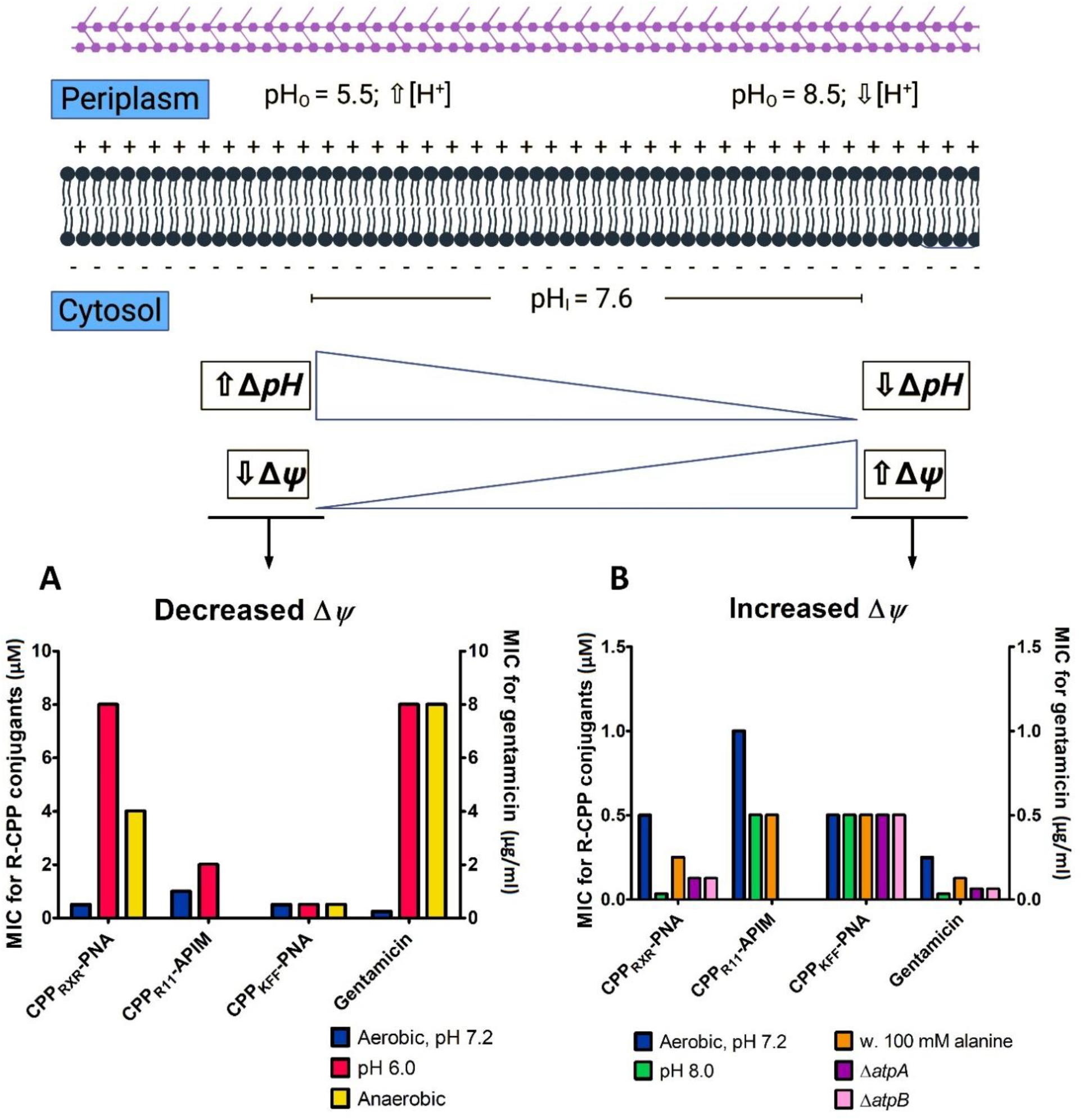
The magnitude of Δ*Ψ* across the cytoplasmic membrane determines CPP_RXR_-PNA and aminoglycoside tolerance. A schematic model of PMF across the cytoplasmic membrane. pH_I_ is pretty constant over the growth pH_o_ range (5.5 to 8.5), but the contributions of Δ*Ψ* and Δ*pH* vary(9). A: MIC determinations for wild-type tested against CPP_RXR_-PNA, CPP_R11_-APIM, CPP_KFF_-PNA and gentamicin anaerobically or at pH 6.0. CPP-PNA/APIM MICs are given in μM (left y-axes) and gentamicin MICs are given in μg/mL (right y-axes). Wild-type MIC are given for aerobic growth conditions at pH 7.2 identical to the growth conditions for the ALE-experiment. B: MIC determinations for wild-type tested against CPP_RXR_-PNA, CPP_R11_-APIM, CPP_KFF_-PNA and gentamicin at pH 8.0, supplemented w. 100 mM alanine, or with either *atpA* or *atpB* deleted. Peptide-PNA MICs are given in μM (left y-axes) and gentamicin MICs are given in μg/mL (right y-axes). Wild-type MIC are given for aerobic growth conditions at pH 7.2 identical to the growth conditions for the ALE-experiment.

### Neither *cpxR_L20Q_* nor the *cpxA_Δ16-17_* mutations conferred resistance to cationic antimicrobial peptides

Cationic antimicrobial peptides represent the biggest class of antimicrobial peptides and the majority of these are amphipathic. Here, Cap11, Cap18, cecropin P1, apidaecin 1B, indolicidin, protamine, and the most well-known polypeptide antibiotic colistin were tested against wild-type, *cpxR_L20Q_, cpxA_Δ16-17_*, and Ahx^R^-3 (Table S4). Colistin, Cap11, Cap18, protamine, and cecropin P1 are believed to target the cell envelope (disruption of bilayers), while indolicidin (disruption of bilayers and DNA synthesis) and apideacin 1B (ribosomes) have intracellular targets. An activated Cpx-response and a decrease in Δ*Ψ* did not confer tolerance to any of the tested AMPs (Table S4). Interestingly, the evolved strain Ahx^R^-3 was more susceptible to Cecropin P1, Cap11, Cap18, and colistin than wild-type (Table S4). One of the major differences between wild-type and Ahx^R^-3 is the two IS1-insertions in *waaB* and *waaO* in Ahx^R^-3 (Table 2). This is expected to result in LPS deficiency, which is associated with an increased susceptibility to cationic antimicrobial peptides in *E. coli*(28). Accordingly, when *waaBO* were deleted in the wild-type (Δ*waaBO*) they became more susceptible to Cecropin P1, Cap11, Cap18, and colistin but not to CPP_R11_-APIM.

### High CPP_RXR_-PNA tolerance is associated with fitness cost

The doubling times of *cpxR_L20Q_* and *cpxA_Δ16-17_* mutants were 28 min and 27 min, respectively, when grown aerobically in MHB I at 37°C and somewhat slower than that of wild-type cells (25 min). The reduced doubling times are likely to result from repression of the respiratory operons in the respective mutants. Of interest, the evolved mutant, Ahx^R^-3 (58 min), which conferred the highest resistance to R-PNA (8 μM vs. 4 μM for the *cpxR_L20Q_* and *cpxA_Δ16-17_* mutants), grew more than twice as slow as both the wild-type and the *cpxR_L20Q_* and *cpxA_Δ16-17_* mutants. This shows that CPP_RXR_-PNA resistance comes with a high fitness cost.

## Discussion

Since the late eighties, natural and synthetic CPPs have been developed and modified for applications in both basic- and applied biology including delivery of therapeutic agents, gene editing and cell imaging. Due to efficient delivery, CPPs as carriers for therapeutic compounds to treat various diseases has gained attraction in recent years(5). Although there are many types of CPPs with different physicochemical properties and applications, R-CPPs, are among the most extensively employed and studied(29). PNA, PMO and AMPs are novel, promising antimicrobial agents, of which PNA and PMO require conjugation to a CPP for delivery to the intracellular targets in bacteria(4). Here, we isolated and characterize resistant strains towards an R-CPP. High-level resistance to the arginine-rich CPP-part was difficult to obtain, came with a high fitness cost, and it conferred cross-resistance to aminoglycosides, which is seen as a clear disadvantage for the use of these carriers in antibacterial therapy.

### The cytoplasmic membrane is the main barrier for CPP_RXR_-PNA

The initial interaction with the envelope differs between CPP_R11_-APIM/CPP_RXR_-PNA and cationic AMPs. Thus, loss of LPS had no effect on the activity of CPP_RXR_-PNA or CPP_R11_-APIM while this sensitized wild-type to the cationic AMPs (Table S4). It therefore remains unclear why LPS deficiency were selected in Ahx^R^-1, 2, and 3. A reduction of Δ*Ψ* in the cytoplasmic membrane was found to diminish/reduce CPP_RXR_-PNA and CPP_R11_-APIM activity, demonstrating that a (large) negative Δ*Ψ* is important for activity, and most likely for providing sufficient free energy required for membrane translocation. The more detailed mechanism of membrane passage is not known, but could rely on direct, local membrane interruption or be aided by one or more still unidentified transporter protein. However, for CPP_RXR_-PNA neither our ALE study nor a previous screening of the KEIO collection(4) provides evidence for the existence of a single CPP_RXR_-PNA transport mechanism. However, we cannot exclude that CPP_RXR_-PNA is the substrate of multiple transport systems, in which case the absence of one would be compensated by the remaining other(s).

### CPP_RXR_-PNA resistance arise in multiple steps

Several genetically different R-PNA^R^ mutants arose both within and across the two lineages in the ALE-experiment. This suggest clonal interference within the populations of each ALE lineage, where mutants will compete not just against the susceptible parent but also against each other(30). Clonal interference influences evolutionary trajectories, which typically leads to a population with one dominant clone(31), as suggested here for clone Ahx^R^-3. The evolutionary paths leading to three genotypes, from the two linages, speaks to either multiple ways to become CPP_RXR_-PNA resistance or that they all, more or less, confer the same universal CPP_RXR_-PNA^R^ phenotype. Nonetheless, resistance development to PNA conjugated to an R-CPP is complex, requiring multiple mutations in independent loci to create the CPP_RXR_-PNA^R^ phenotype (Table 2). This is in stark contrast to the straightforward resistance development to PNA conjugated to the lysine-rich CPP, i.e. CPP_KFF_-PNA, by a loss-of-function mutation in the *E. coli sbmA* gene(4). We show that the *cpx*\*-phenotype is a major part of the CPP_RXR_-PNA resistance phenotype in Ahx^R^-3, but that additional mutations are required for high-level resistance. As observed for *waaBO*, not all mutations from the ALE-experiment on their own resulted in CPP_RXR_-PNA tolerance. Either, the *waaBO* mutation requires another mutation to synergistically provide CPP_RXR_-PNA tolerance or it was selected as a mutation that compensate a fitness cost associated with mutations conferring CPP_RXR_-PNA resistance. Indeed, CPP_RXR_-PNA^R^ strains have increased doubling time, highlighting that CPP_RXR_-PNA resistance comes with a cost.

### The cpx* phenotype relies on a functional Cpx-system

Our results show that alterations in the first transmembrane domain of CpxA can produce a *cpx*\* phenotype. Several gain-of-function mutations in *cpxA* have been reported(16, 17), but most cluster to and around the second transmembrane domain of CpxA(17). We suggest that the deletion in the first transmembrane domain leads to a conformational change in the periplasmic part of CpxA, which either disallows CpxP regulation or simulates the stress signal that actives the Cpx-response; i.e. a constitutive active CpxA. To the best of our knowledge, *cpxR_L20Q_* is the first reported mutation in *cpxR* resulting in a *cpx*\* phenotype. The activated Cpx-response in the *cpxR_L20Q_* mutant was CpxA-dependent; i.e. the *cpxR_L20Q_* mutation does not result in a constitutive phosphorylated CpxR. Therefore, it is likely that the *cpxR_L20Q_* mutation inhibits/diminish the ability of CpxA to dephosphorylate CpxR-P to CpxR, which in turn results in a constitutive active Cpx-response. CpxA and CpxR-P both function as dimers(32). Thus overproducing the respective wild-type allele in the two mutants restored sensitivity to CPP_RXR_-PNA by either hetero-dimerization of a mutated and a wild-type protein, with the wild-type being dominant to the mutant, or by homo-dimerization of either mutated or wild-type proteins, but with wild-type dimer domination due to abundance.

### Low cytoplasmic membrane potential correlates with tolerance to CPP_RXR_-PNA, CPP_R11_-APIM, and aminoglycosides

We found that a reduction of the Δ*Ψ* component of PMF across the cytoplasmic membrane increased tolerance to both CPP_RXR_-PNA, CPP_R11_-APIM, and aminoglycosides; the latter in agreement with previous studies in *Bacillus subtilis* and *E. coli*(33). The reduced activity of both CPP_RXR_-PNA, CPP_R11_-APIM, and aminoglycosides when Δ*Ψ* was decreased indicates that crossing of the inner (and not the outer) membrane is the rate limiting and energy dependent step shared by this type of (polycationic) compounds as previously proposed for aminoglycosides (34). Once inside the bacterial cells, aminoglycosides interacts with the 30S subunit of ribosomes, RXR-PNA prevent *acpP* translation, and CPP_R11_-APIM prevent β-clamp function (Fig. 4A). In wild-type cells, CPP_RXR_-PNA (and possibly also aminoglycosides and, CPP_R11_-APIM) do not activate the Cpx-response. We propose that *cpx*\* dependent tolerance to CPP_RXR_-PNA, CPP_R11_-APIM, and aminoglycosides is caused by downregulation of respiratory complexes in the cytoplasmic membrane (Fig. 4B) thereby decreasing Δ*Ψ* across the cytoplasmic membrane at least partly due to the reduced ATP level and consequently reduced proton chemical potential over the membrane, making it less energetically favorable for R-CPP to transvers the cytoplasmic membrane (Fig. 4B). CPP_KFF_-PNA is internalized in a SbmA-dependent manner(4) (for which a membrane potential driving force apparently is not important) and therefore the CPP_RXR_-PNA^R^ strains were sensitive to CPP_KFF_-PNA.

**Fig. 4.**
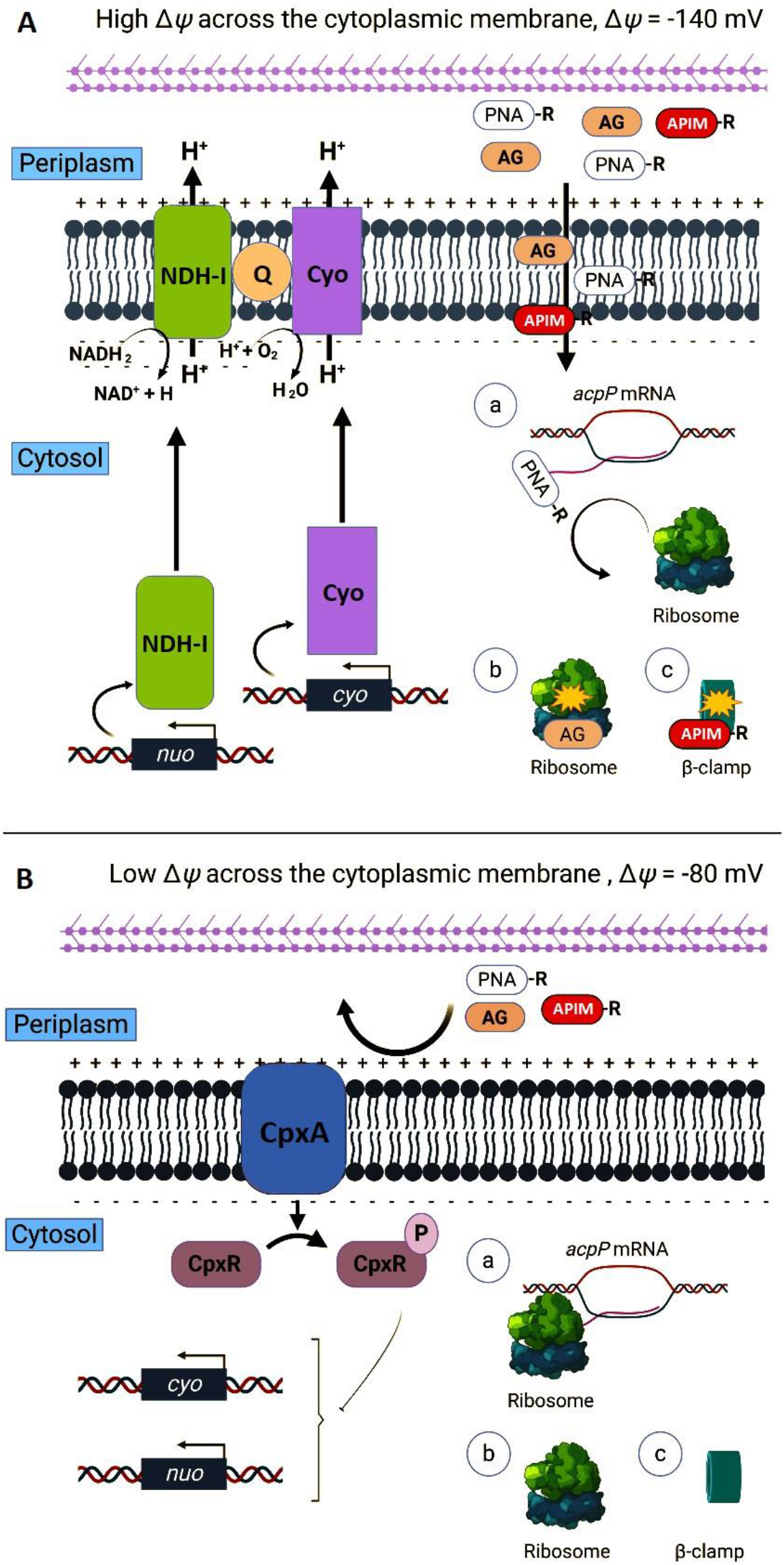
CPP_RXR_-PNA, CPP_R11_-APIM, and aminoglycoside uptake wild-type (A) and *cpx*\* (B). A schematic model of CPP_RXR_-PNA, CPP_R11_-APIM, and aminoglycoside (AG) internalization across the cytoplasmic membrane and action in wild-type (A) and a *cpx*\*mutant (B). A: In the wild-type CPP_RXR_-PNA, CPP_R11_-APIM, and AG crossing of the cytoplasmic membrane from the periplasmic space and into the cytosol correlates with the magnitude of the Δ*Ψ*. The respiratory complexes NADH:ubiquinone oxidoreductase complex I (NDH-I) transcribed from the *nuoABCEFGHIJKLMN*-operon (*nuo*) and cytochrome *bo*_3_ ubiquinol oxidase (Cyo) transcribed from the *cyoABCDE* operon (*cyo*) are shown. CPP_RXR_-PNA, AG, and CPP_R11_-APIM in the cytosol leads to growth arrest by steric hindrance of ribosome binding to *acpP* mRNA for CPP_RXR_-PNA (A.a), binding to the 30S subunit of ribosomes impairing translational accuracy (A.b), and binding to the β-clamp, respectively. B: In a *cpx*\* mutant CPP_RXR_-PNA, CPP_R11_-APIM, and AG are not able to transvers the cytoplasmic membrane from the periplasmic space and into the cytosol. The constitutive active Cpx-response downregulates *nuo, cyo*, which results in a reduced Δ*Ψ*, which correlates with reducing killing for all the compounds. Lack of CPP_RXR_-PNA and AG in the cytosol leads to translation of *acpP* mRNA (B.a) wild-type like 30S ribosomal function (B.b), and wild-type like β-clamp function, respectively, both leading to survival during CPP_RXR_-PNA, CPP_R11_-APIM, and AG treatment.

### The cpx* phenotype could lead to the acquisition of other mutations to promote resistance or persistance

The *cpx*\* phenotype provides moderate resistance to both CPP_RXR_-PNA and aminoglycosides – all without a major loss of fitness. Thus, in a clinical setting, *cpx*\* mutants could grow in the presence of moderate levels of either compound, which in turn would create an evolutionary opportunity to adapt and evolve by subsequent acquisition of mutations to further increase resistance or to assist in persistent colonization. Several recent studies have shown that adaptation, may have a significant impact on the selection of highly resistant/persistent clones for *Mycobacterium* (ribosomal mutations)(35), *Pseudomonas aeruginosa* (efflux pump regulator mutation)(36), and *E. coli* (formation of subpopulations with an increased overall mutation rate)(37). Thus, the *cpx*\* phenotype may be added to this list.

## Materials and Methods

### Growth conditions

Cells were grown in Luria–Bertani Broth (LB) medium or Müller-Hinton I Broth (MHBI) at 37°C with aeration unless stated otherwise. When necessary, antibiotics were added to the following concentrations: chloramphenicol, 20 μg/mL; ampicillin, 150 μg/mL; kanamycin, 50 μg/mL; tetracyclin, 10 *μ*g/mL.

### Peptide conjugates

PNA-peptide conjugates were synthezised as described previously (1, 4). CPP_R11_-APIM (RWLVK* with the complete sequence Ac-MD-RWLVK-GILQWRKI-RRRRRRRRRRR) is described previously(22).

### Bacterial strains and plasmids

A MG1655 strains used in this study are listed in Table S5 and all primer sequences in Table S6. For deleting and overexpressing genes, we used the KEIO collection(38) and the ASKA library(39), respectively. Shortly, the *cpxA, cpxR, degP*, or *cpxP* deletion/replacement mutations were obtained from the KEIO collection(38) and P1 transduced(40) into *E. coli* MG1655 to create ALO6524, ALO6525, ALO7548, and ALO7549, respectively. For overexpression of genes, pCA24n based plasmids from the ASKA collection was transformed into *E. coli* MG1655 selecting for chloramphenicol.

Deletion of the *nuoABCEFGHIJKLMN, cyoABCDE*, and *waaBO* chromosomal regions in wild-type was constructed by the lambda red procedure(41). Briefly, primer sequences were obtained from Baba *et al*. (38) and are listed in Suppl. Table 2. The *nuo*_cat_Fw primer corresponds to the 5’ primer used to delete *nuoA* in Baba *et al*. (38), while the *nuo*_cat_Rv primer corresponds to the 3’ primer used to delete *nuoN* in Baba *et al*. (38). The *cyo*_cat_Fw primer corresponds to the 5’ primer used to delete *cyoA* in Baba *et al*. (38), while the *cyo*_cat_Rv primer corresponds to the 3’ primer used to delete *cyoE* in Baba *et al*. (38). The *waaB*_cat_Fw primer corresponds to the 5’ primer used to delete *waaB* in Baba *et al*. (38), while the *waaO*_cat_Rv primer corresponds to the 3’ primer used to delete *waaO* in Baba *et al*. (38). DNA fragments were PCR amplified using the primer sets *nuo*_cat_Fw + *nuo*_cat_Rv, *cyo*_cat_Fw + *cyo*_cat_Rv, and *waaB*_cat_Fw + *waaO*_cat_Rv to delete *nuoABCEFGHIJKLMN, cyoABCDE*, and *waaBO*, respectively. PCR amplification was done using high-fidelity Phusion DNA polymerase (Thermo Scientific™) according to the manufacturer’s specifications with pKD3 as template. The resultant DNA fragments were introduced into wild-type cells bearing pKD46(41), selecting for chloramphenicol resistance. Each deletion was verified by PCR creating Δ*nuoABCEFGHIJKLMN* (Δ*nuo*; ALO7555), Δ*cyoABCDE* (Δ*cyo*; ALO7556), and Δ*waaBO* (ALO6972).

The *cpxR_L20Q_* mutations were introduced into the wild-type using the Tet/SacB counter-selection and recombineering approach(42). Briefly, the *tet-sacB* cassette flanked by appropriate homology arms was amplified using the primer set *CpxR_tet-sacB_Fw* + *CpxR_tet-sacB_Rv* using pTETSACB as a template for exchange of *cpxR* in the wild-type. The resultant DNA fragments were introduced into wild-type bearing pKD46(41), selecting for tetracycline and ampicillin resistance at 32°C (to keep pKD46 in the cell) creating *cpxR:tet-sacB* pKD46. The deletion was verified by PCR. Next, the cpxR_L20Q_ mutations were amplified from Ahx^R^-3 with homology flaking the inserted *tet-sacB* cassettes, using the primer sets cpxR_L20Q__Fw + cpxR_L20Q__Rv. The resultant DNA fragments containing the *cpxR_L20Q_* mutation were introduced into *cpxR:tet-sacB* selecting for growth on LB agar containing 6% sucrose. *cpxR* genes from clones that were tetracycline sensitive and able to grow on sucrose were sequenced to confirm the introduction of the specific mutations from Ahx^R^-3 into wild-type, thus creating the strain *cpxR_L20Q_* (ALO7180). *cpxA* was deleted in *cpxR_L20Q_* and *cpxR* was deleted in *cpxA_Δ16-17_* by P1 transduction (as above) creating *cpxR_L20Q_ ΔcpxA* (ALO7550) and *cpxA_Δ16-17_ ΔcpxR* (ALO7551), respectively. In addition, *nuoABCEFGHIJKLMN* and *cyoABCDE* were deleted as above in *cpxR_L20Q_* using the lambda red procedure(41) created *cpxR_L20Q_* Δ*nuo* (ALO7552) and *cpxR_L20Q_* Δ*cyo* (ALO7553), respectively.

### Adaptive Laboratory Evolution (ALE) experiment

Wild-type was evolved for 20 days to CPP_RXR_-PNA, with two lineages evolved in parallel for each drug. Here, 1 x 10^5^ CFU/mL of cells were inoculated in 2-fold peptide-PNA gradient in 8 dilutions in 100 μl MHBI in a 96-well polystyrene microtitre plate and grown for 18-24 hours at 37°C. The following day the population grown in the highest concentration of peptide-PNA where re-diluted to 1 x 10^5^ CFU/mL and inoculated in a fresh 2-fold peptide-PNA gradient as above. After 20 successive passages, five peptide-PNA resistant single clones were isolated from each lineage and whole genome sequenced. As control, *E. coli* MG1655 was evolved to the MHBI in four independent biological replicates without the presence of peptide-PNA. Here, two clones were isolated and whole genome sequenced for each lineage.

### Extraction of genomic DNA

Genomic DNA of *E. coli* MG1655 was isolated using phenol/chloroform extraction. Shortly, cells from a 15 ml culture were collected at an OD600 of 0.5 by centrifugation at 5.000 ×g for 10 minutes. The pellet was resuspeded in 1 ml 0.9% NaCl, followed by a centrifugation at 10.000 ×g for 5 minutes. The cells were lysed by incubation at 37°C for 1 hour with 600 μl lysis buffer (Tris-EDTA buffer (TE buffer, pH 8.0), 10% (w/v) SDS, 20 mg/ml proteinase K, 1 mg/ml RNase A). Following two phenol- and two chloroform-extraction, DNA was precipitated using NaCl (0.2 M) and 0.6 vol. isopropanol, washed in 80% ethanol and resuspended in TE buffer (pH 8.0).

### Whole genome sequencing and data analysis

Reads were mapped to MG1655 NC_000913.3 using BWA-MEM. Variants were extracted using Freebayes (only variants with >50 % frequency were retained). For location of IS insertion, mapped reads with <60 mapQuality were selected and paired reads (CIGAR-Left-clip/Right-Clip, CIGAR-D and CIGAR-I) were aligned to NC_000913.3 using NCBI Blast to confirm deletion /insertion.

### Antimicrobial Susceptibility Testing

MIC values were determined by broth microdilution according to the standard protocol(43) with a few modifications. Shorty, an overnight bacterial cell culture was diluted to approximately 5 × 10^5^ CFU/mL in MHBI. 100 μL bacterial solution was dispensed into a low-bind 96-well plate (Thermo-Scientific, cat.no. 260895) along with 11 μL of the test compound (2-fold dilutions). The plate was incubated at 37°C for 18-24 h. The MIC was determined as the lowest concentration, which inhibited visible growth in the wells (OD_595_ nm < 0.1). Peptide-PNAs were dissolved in 0,02% acetic acid containing 0,4% bovine serum albumine and dilution were performed in 0,01% acetic acid containing 0,2% BSA, while gentamicin, amikacin, kanamycin, cecropin P1, Cap11 (Genescript), Cap18 (Genescript), apidaecin 1B (Genescript), colistin, indolicidin and protamine (Sigma-Aldrich) was diluted according to vendors specifications. To determine MIC under acidic and alkaline conditions, MHBI was buffered to pH 6.0 or pH 8.0 using citrate buffer (pH 3.0) and carbonate-bicarbonate buffer (pH 10.0), respectively. Anaerobic growth for MIC determinations was performed in a double sealed bag using anaerobic atmosphere generation bags (Becton, Dickinson and Company, New Jersey, USA). MHBI was supplemented with 0.1 mM Isopropyl β-D-1-thiogalactopyranoside (IPTG), when tested strains harbored pCA24n-based plasmids.

### Total RNA extraction

RNA were isolated from cells by the hot phenol method. Shortly, cells were grown in cation adjusted MHBI to mid-log phase, an OD_595_ of 0.2 (2.5 × 10^9^ cells/ml), by shaking at 150 rpm and 37°C. To induce protein expression in cells harboring the inducible pCA24n-based plasmid, IPTG was added to a concentration of 0.1 mM at an optical density OD_595_ 0.005. Bacteria from 2-ml aliquot were harvested by centrifugation at 4°C and 17.000×g for 5 minutes. The pellets were resuspended in the same volume of RNAprotect Bacteria Reagent (QIAGEN, Denmark), incubated at room temperature for 5 minutes, followed by a centrifugation using the same conditions as above. The supernatants were discarded and the pellets were either stored at −80°C until needed or immediately used for RNA extraction.

To extract RNA, the pellets were lyzed in 200 μl Solution I [0.3 M sucrose, 0.01 M sodium acetate (NaOAc) and pH 4.5] at 4°C and mixed thoroughly. Then, 200 μl of Solution II (2% (w/w) Sodium dodecyl sulfate (SDS), 0.01 M NaOAc and pH 4.5) were added at room temperature and solutions were gently mixed by inversion of tubes. Next, a volume of 400 μl of acid phenol:chloroform:isoamyl alcohol 125:24:1 was added and immediately mixed (vortex) to emulsify. The samples were incubated at 65°C for 10 minutes and mixed (vortex) for 5-10 seconds every minute. Then, tubes were transferred on ice for 5 minutes and centrifuged at 14.000 rpm for 5 minutes. The aqueous phase was transferred to a new polypropylene tube and one-fifth volume of chloroform was added and gently mixed by inversion, followed by centrifugation at 14.000 rpm for 5 minutes. This step was repeated by addition of the same volume of chloroform:isoamyl alcohol 24:1. To precipitate RNA, the aqueous phase was transferred to a new tube, mixed with the same volume of isopropanol and incubated at −80°C for a minimum of 1 hour. RNA was precipitated by spinning down at 4°C and 12.000 ×g for 30 minutes, followed by wash with 1 ml 70% ethanol and centrifugation at 4°C and 12.000 ×g for 5 minutes. The pellets were resuspended in 30 μl nuclease free water and RNA concentration and purity was determined using a NanoDrop.

Following removal of genomic DNA by DNase I (Qiagen, Venlo, Netherlands), RNA were purified using RNeasy MinElute Cleanup kit (Qiagen) and following the vendors instructions.

### qRT-PCR

For qRT-PCR experiments, after genomic DNA digestion, 1 μg of total RNA was retrotranscribed using QuantiTect Reverse Transcription kit (Qiagen). Primers designed to amplify *cpxP*, *nuoA*, and *cyoA* (Table 8) were targeted to regions of unique sequences within the genes in wild-type *E. coli* MG1655. The qRT-PCR was performed using the TB Green Premix Ex Taq II (Takara Bio Inc., Shiga, Japan) on a QuantStudio 3 Real-Time PCR System (Thermo Fisher Scientific, Massachusetts, United States). All data were normalized to the reference genes *hctA* and *cysG*. These data were transformed to log2 to obtain a change difference (n-fold) between strains.

### Determination of membrane potential (Δ*ψ*) by measuring Tetraphenyl phosphonium (TTP^+^) uptake using the TPP^+^-selective electrode method

The membrane potential was determined by measuring the uptake of the permeating lipophilic cationic probe, Tetraphenyl phosphonium (Merck KGaA, Germany). An overnight culture was diluted (5.000-fold) in MHBI, and cells from 25 ml culture were harvested when OD_595_ reached 0.2. Cells were washed twice in 0.1 M Tris·HCl (pH 8.1) using the conditions mentioned above, and the pellets were gently resuspended in 1 ml of the same buffer. To generate cells permeable to TPP^+^, after a first incubation for 6 minutes at 36°C with occasional agitation, bacteria were treated with K^+^-EDTA to a concentration of 10 mM and then incubated at 36°C for 3 more minutes. Cells were diluted 10-fold in ice-cold 0.1 M potassium phosphate buffer (0.1 M K_2_HPO_4_; 0.1 M KH_2_PO_4_; pH 7.5), to remove EDTA, followed by immediate centrifugation in the cold at 13.000 rpm for 7 minutes. The cells were washed twice in the same ice-cold buffer using the parameters mentioned above, and the pellets were resuspended, to an OD_595_ of 20.0, in 10 mM potassium phosphate buffer (pH 7.5) containing 200 mM NaCl, 0.14 mM CaCl_2_, 0.1 mM MgSO_4_ and 0.01 mM MnCl_2_. The cells were kept on ice until use.

Bacteria were added to an OD_595_ of 4.0 in 10 ml of the same buffer supplemented with glucose and TPP^+^ to a concentration of 5 mM and 10 μM, respectively, and TPP^+^ uptake was measured utilizing the TPP^+^-selective electrode method described by Hosoi *et al*(44). TPP^+^ concentration in the external medium was determined using a Kwik-Tip^TM^ Ag/AgCl half-cell electrode, which was constructed according to the instructions provided by the manufacturer (World Precision Instruments, Sarasota, FL). Both the TPP^+^-selective electrode and a Flexible Dri-Ref reference electrode (World Precision Instruments, Sarasota, FL) were connected to an Jenway 3510 ion meter (Cole-Parmer, Staffordshire, UK) and LabTrax 4-Channel Data Acquisition system (WPI). Finally, TPP^+^ uptake measurements were carried out in a 15 ml closed tube, at 30°C and pH 7.5 for 10 minutes, and data were recorded using the iWorx LabScribe recording and analysis software (iWorx Systems Inc., Dover, NH).

## Acknowledgments

This paper is dedicated to the memory of Mads Christian Guldbæk (1991-2019).

## Author contributions

Conceptualization: JFM, PEN, and ALO; data curation: JFM and AK; formal analysis: JFM, AK, and ALO; investigation: JFM and AK; methodology: JFM, AK, GC, and ALO; project administration: JFM and ALO; resources: JFM, PEN, and ALO; bioinfomatics: GC; supervision: PEN and ALO; manuscript writing: JFM; manuscript editing: JFM, AK, GC, MO, PEN, and ALO.

## Data and materials availability

All data needed to evaluate the conclusions in the paper are present in the paper and/or the Supplementary Materials. Additional data are available from authors upon request. All figures were created with BioRender.com.

## Supplementary Materials

**Table S1.**
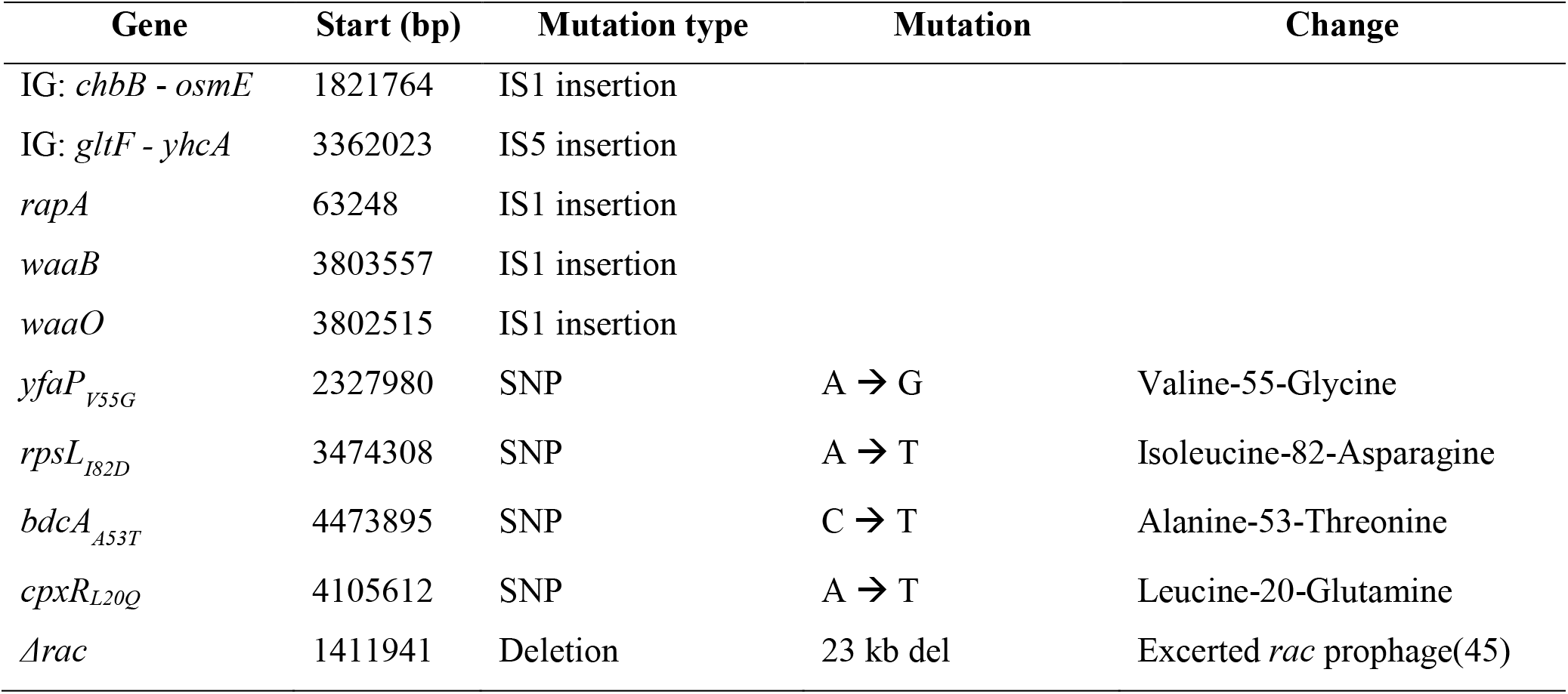
Mutation type.

**Table S2.** Function of mutated genes.

| Gene | Function | Reference |
| --- | --- | --- |
| *chbB* | *N,N'*-diacetylchitobiose-specific phosphotransferase system enzyme IIB component | (46) |
| *osmE* | Osmotically-inducible lipoprotein, which is induced at the onset of stationary phase | (47) |
| *gltF* | Unknown | (18) |
| *yhcA* | Hypothetical protein, function unknown |  |
| *rapA* | RapA enables recycling of stalled RNA polymerases and stimulates RNA synthesis *in vitro* | (48) |
| *waaB* | The *waa*-operon consists of *waaQGPSBOJYZU* whose products are involved in the assembly of the core region of the lipopolysaccharide molecule in the outer membrane of the cell envelope. | (49) |
| *waaO* |  |  |
| *yfaP* | Unknown |  |
| *rpsL* | The product of *rpsL* is the S12 protein; a component of the 30S subunit of the ribosome that plays a role in translational accuracy. | (20) |
| *bdcA* | BdcA is a c-di-GMP-binding biofilm dispersal mediator protein | (50) |
| *cpxA* | *cpxAR* encodes the two-component regulatory system CpxA/R, that consists of the transmembrane sensor kinase CpxA in conjunction with the cytoplasmic transcription factor CpxR. | (21) |
| *cpxR* |  |  |

**Table S3.** Accumulated fitness gain mutations from the ALE-experiment to the media alone.

| Gene | Start (bp) | Mutation type | Mutation | Change |
| --- | --- | --- | --- | --- |
| *yiaD* | 3716565 | SNP | C → T | Leucine-7-Phenylalanine |
| *uspC* | 1979381 | IS1 insertion |  |  |
| *uspC* | 1979385 | IS1 insertion |  |  |
| *uspC* | 1979486 | IS1 insertion |  |  |
| *uspC* | 1979489 | IS1 insertion |  |  |
| *uspC* | 1979507 | IS1 insertion |  |  |
| *uspC* | 1979510 | IS1 insertion |  |  |
| *ccmA* | 2297378 | IS1 insertion |  |  |
| *ccmA* | 2297389 | IS1 insertion |  |  |
| *fimE* | 4542161 | Inversion/duplication |  |  |
| *fimE* | 4542164 | Inversion/duplication |  |  |
| *fimE* | 4542682 | Inversion/duplication |  |  |
| *fimE* | 4542689 | Inversion/duplication |  |  |
| *fimE* | 4542690 | Inversion/duplication |  |  |
| *fimE* | 4542987 | Inversion/duplication |  |  |
| *fimE* | 4542995 | Inversion/duplication |  |  |

**Table S4.** cpxR_L20Q_ and cpxA_Α16-17_ does not provide tolerance to cationic antimicrobial peptides.

MIC values for peptide-PNA (in μM) and AMPs (in μg/mL) for wild-type and indicated mutants. MIC were determined using broth dilutions (see Methods) at 37°C (no shaking). Peptide-PNA sequences are listed in Table 1.
| Minimum inhibitory concentration to peptide-PNA/APIM (in μM) and AMPs (in μg/mL) |  |  |  |  |  |
| --- | --- | --- | --- | --- | --- |
| AMP/strain | Wild-type | *cpxA<sub>Δ16-17</sub>* | *cpxR<sub>L20Q</sub>* | *ΔwaaBO* | Ahx<sup>R</sup>-3 |
| CPP<sub>RXR</sub>-PNA | 0.5 | 4 | 4 | 0.5 | 8 |
| CPP<sub>R11</sub>-APIM | 1 | 2 | 2 | 1 | N/A |
| Cecropin P1 | 32 | 32 | 32 | 16 | 16 |
| Cap11 | 2 | 2 | 2 | 1 | 1 |
| Cap18 | 2 | 2 | 2 | 1 | 1 |
| Indolicidin | 4 | 4 | 4 | 4 | 4 |
| Apidaecin 1B | 8 | 8 | 8 | 8 | 8 |
| Colistin | 0.125 | 0.125 | 0.125 | 0.0625 | 0.0625 |
| Protamine | 16 | 16 | 16 | 16 | 16 |

**Table S5.**
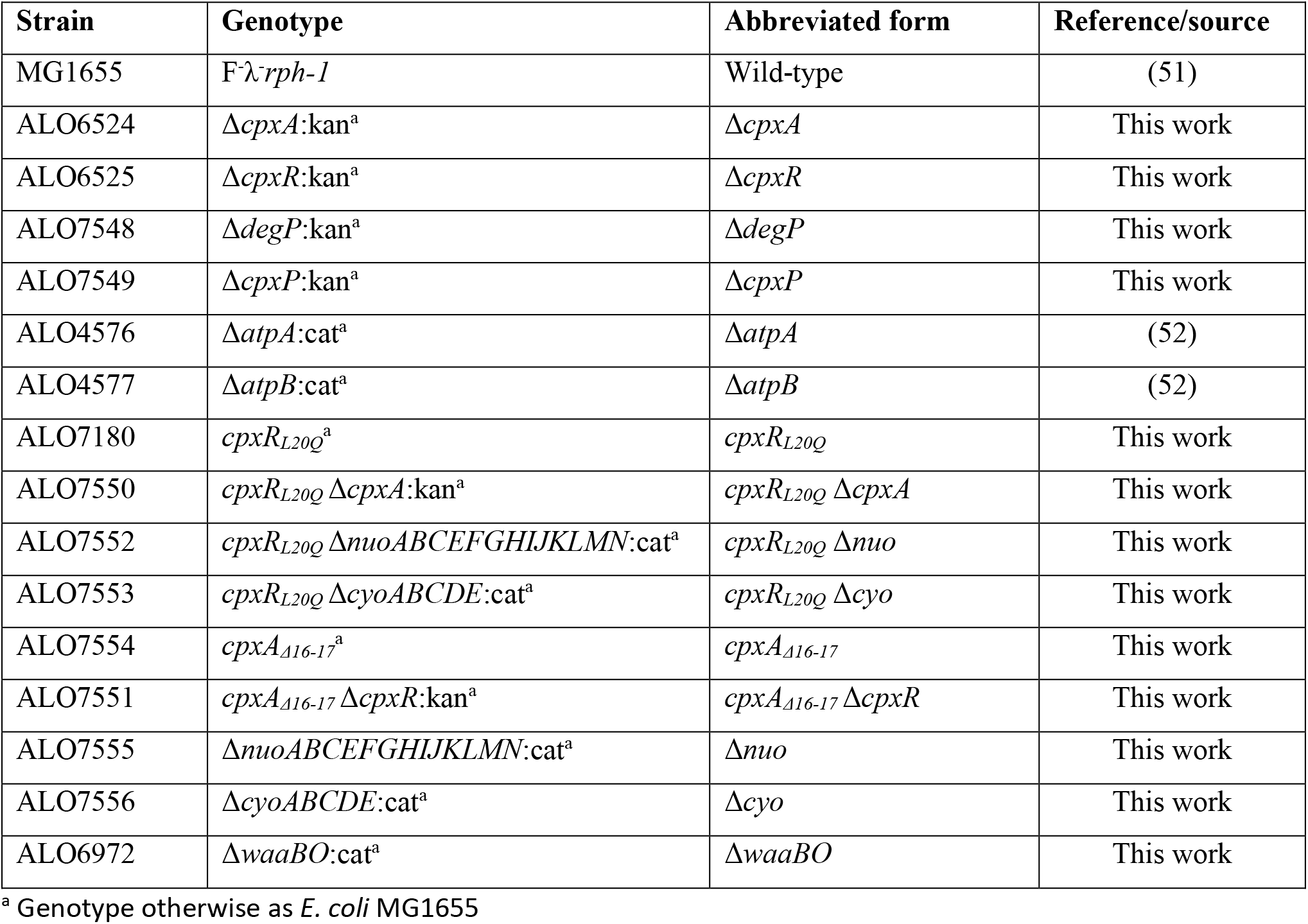
Bacterial strains.

**Table S6.**
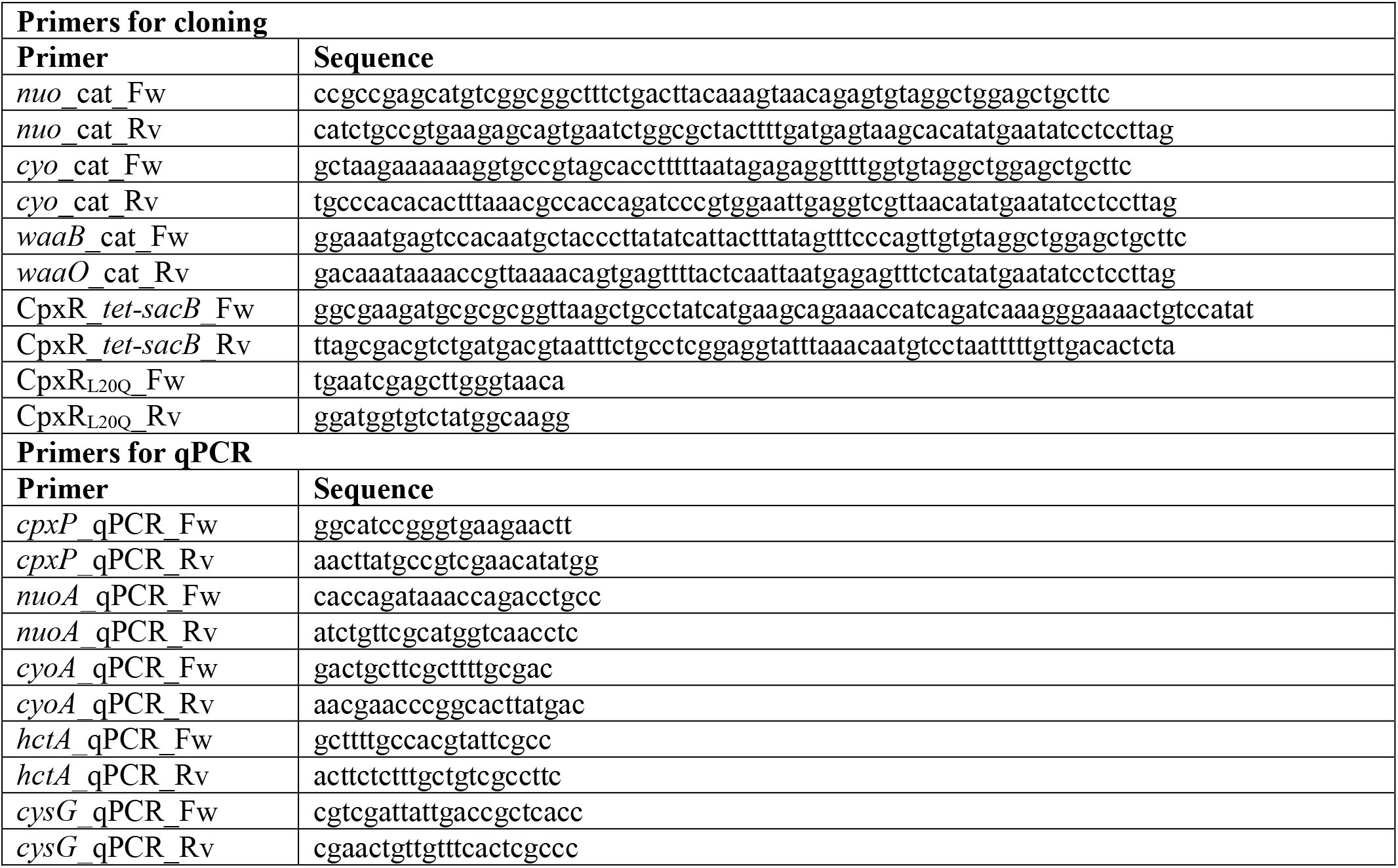
Primer sequences.

